# Cleanliness in context: reconciling hygiene with a modern microbial perspective

**DOI:** 10.1101/095745

**Authors:** Roo Vandegrift, Ashley C. Bateman, Kyla N. Siemens, May Nguyen, Jessica L. Green, Kevin G. Van Den Wymelenberg, Roxana J. Hickey

**Affiliations:** Biology and the Built Environment Center, University of Oregon; Institute of Ecology and Evolution, Department of Biological Sciences, University of Oregon; Energy Studies in Buildings Laboratory, Department of Architecture, University of Oregon

**Keywords:** Hygiene, microbiota, microbiome, skin, microbial ecology, hand drying

## Abstract

The concept of hygiene is rooted in the relationship between cleanliness and the maintenance of good health. Since the widespread acceptance of the germ theory of disease, hygiene has become increasingly conflated with that of sterilization. Recent research on microbial ecology is demonstrating that humans have intimate and evolutionarily significant relationships with a diverse assemblage of microorganisms (our *microbiota*). Human skin is home to a diverse, skin habitat specific community of microorganisms; this includes members that exist across the ecological spectrum from pathogen through commensal to mutualist. Most evidence suggests that the skin microbiota is likely of direct benefit to the host, and only rarely exhibits pathogenicity. This complex ecological context suggests that the conception of hygiene as a unilateral reduction or removal of microbes has outlived its usefulness. As such, we suggest the explicit definition of hygiene as ‘**those actions and practices that reduce the spread or transmission of pathogenic microorganisms, and thus reduce the incidence of disease**’. To examine the implications of this definition, we review the literature related to hand drying as an aspect of hand hygienic practice. Research on hand drying generally focuses on ‘hygienic efficacy’, a concept not typically defined explicitly, but nearly always including alterations to bulk microbial load. The corresponding literature is differentiable into two divisions: research supporting the use of forced air dryers, which typically includes effectiveness of drying as an aspect of hygienic efficacy; and research supporting the use of paper towels, which typically includes risk of aerosolized spread of microbes from hands as an aspect of hygienic efficacy. Utilizing a definition of hygiene that explicitly relies on *reduction in disease spread* rather than alterations to bulk microbial load would address concerns raised on both sides of the debate. Future research should take advantage of cultivation-independent techniques, working to bridge the gap between the two existing divisions of research by using health outcomes (such as the spread of disease) as dependent variables, taking into account the microbial community context of the skin microbiota, and focusing on understanding the relative contribution of bioaerosols and residual moisture to the risk of disease transmission.

## Background

This Review focuses on the concept of hygiene as it relates to the human-associated microbiota, with the aim of coming to a clear, workable definition of hygiene that is congruent with our emerging understanding of the intimate, multifaceted, and symbiotic relationships that humans have with microorganisms. We examine clinical and commonplace definitions of hygiene and re-evaluate the concept in the context of a modern understanding of human-associated microbial ecology, both *on* humans and *around* humans in the built environment (BE). By doing this, we bridge the gap between the clinical skin microbiology literature and the emerging human-associated microbial ecology literature. Our Review closes with a targeted analysis of a specific segment of scientific literature relevant to public health: the body of work on *hand drying* as an aspect of hand hygiene. We use hand drying as a case-study to examine the implications of using a microbial ecology-based approach to defining hygiene.

The word *hygiene* originates with Hygieia, the Greek goddess of health. The Oxford English Dictionary (OED) defines it thus: “That department of knowledge or practice which relates to the maintenance of health; a system of principles or rules for preserving or promoting health; sanitary science” [1] and the Merriam-Webster defines it as: “a science of the establishment and maintenance of health; conditions or practices (as of cleanliness) conducive to health” [2]. The OED also gives us some context of the use of the word in English, noting that its origins lay with the first part of the definition: early use of the word relates entirely to the practice of medicine (e.g., in 1671 W. Salmon writes that medicine has three parts — physiology, hygiene, and pathology). More modern usage, however, has shifted to the latter definition; hygiene in most modern contexts tends to refer specifically to the practice of cleanliness where it relates to maintaining good health (e.g., “dental hygiene” referring to the practice of regular cleaning of the teeth). In practice, however, hygiene is rarely explicitly defined. The term most often refers to *hand hygiene*, which the World Health Organization defines as “a general term referring to any action of hand cleansing” [3]. Hygiene may also refer to *environmental hygiene*, which can mean either the cleaning of surfaces within a person’s (most commonly a patient’s) environment [4], or, more broadly, infrastructural changes that alter the environment in a way perceived as beneficial to human health (such as the installation of water and sewage treatment facilities) [5].

We focus primarily on hand hygiene, since this aspect of hygiene is most commonly used in the modern scientific literature, and draw parallels with environmental hygiene in light of recent developments in indoor microbial ecology.

### History and regulation of hand hygiene

Interest in hand hygiene dates to the middle of the 19th century. Oliver Wendell Holmes (1809 – 1894), in Boston, and Ignaz Philipp Semmelweis (1818 – 1865), in Vienna, both noticed the contagious nature of puerperal fever, which affects women shortly after childbirth [6,7]. Publishing their findings concurrently, but on different continents, they both argued that physicians with unwashed hands spread the disease to birthing women. Semmelweis’s work went one step further; he made the connection that medical students often went straight from the autopsy theatre to the birthing room, and concluded that they must be transmitting “cadaverous particles” from the corpses to the patients. To combat this spread, he instituted a policy of scrubbing the hands in chloride of lime (calcium hypochlorite) for anyone moving between the autopsy theatre and the maternity wards; mortality rates were quickly reduced [8].

Both physicians were ridiculed for their beliefs at the time, but they laid the foundations for thought about hygiene and the spread of infection in the medical establishment. Around this time, in France, Louis Pasteur was working on germ theory and fermentation, formally publishing the pasteurization method in 1865, followed by the initial publication on germ theory in silkworms in 1870, just nine years after Semmelweis’s research on puerperal fever [9]. Pasteur was also working on puerperal fever; in 1880, he published microbiological observation and recommendations concerning the disease [10], which were more readily accepted by the medical establishment than Semmelweis’s recommendations.

Despite this early recognition of the importance of hand hygiene, little attention was paid to the particulars for most of the next century. In the United States, regulation started with the issuance of formal recommendations by the US Public Health Service in 1961 [11] that healthcare workers (HCWs) must wash their hands with soap and water for at least one minute before and after patient contact. From there, increasing recognition of the importance of hand hygiene in infection control and prevention, particularly in hospital and other healthcare-related settings, led to the publication of guidelines and recommendations by the CDC in 1975 and 1985 [12,13]. Recognition of the spread of antibiotic resistance in hospitals, along with the role of low compliance to existing hand hygiene policies, prompted further regulation [14,15], leading eventually to new regulations from the CDC in 2002 [16], including recommended use of alcohol rubs in place of handwashing for the first time as a means of increasing compliance with hand hygiene recommendations. This encouraged the widespread adoption of alcohol rubs around the world [17]. Internationally, regulation culminates in the World Health Organization’s (WHO) First Global Patient Safety Challenge in 2005, which focuses on hand hygiene and led to the eventual publication of the comprehensive *Guidelines on Hand Hygiene* in 2009 [3], which now serves as the international standard for hand hygiene practices.

The WHO explicitly defines hand hygiene as “any action of hand cleansing”, and then goes on to delineate many specific “hand hygiene practices”, which include everything from soap and water handwashing to surgical hand antisepsis. It is noteworthy that **most regulations and recommendations concerning hand hygiene focus on the aspect of hygiene as the act of cleaning, concentrating on the reduction in bulk microbial load, rather than reduction in transmission of infection.**

### Environmental hygiene

In addition to the focus on the importance of hand hygiene to the prevention of healthcare-associated or nosocomial infection, there is growing recognition that other aspects of hygiene beyond hand hygiene are also important [4,18]. These are often collectively referred to as *environmental hygiene*, which can be further classified into two distinct groups: community environmental hygiene and environmental hygiene of the built environment.

Community environmental hygiene refers to those aspects of hygiene that affect entire communities of people, and typically involves infrastructural issues such as the availability of potable water and sanitary disposal of human wastes. The effects of improvements of community environmental hygiene are relatively well established [3,19–22], and are generally accepted to be of great importance to the reduction in transmission of certain diseases, particularly those with a fecal-oral transmission route [5]. The WHO Global Water Supply and Sanitation Assessment identifies three key hygiene behaviors most likely to improve community health: (1) handwashing, (2) safe disposal of feces (particularly those of children), and (3) safe water handling and storage [5].

Environmental hygiene of the built environment (EHBE) typically refers to practices of cleaning surfaces for the purposes of reducing transmission of pathogens via those surfaces (e.g., liquid disinfectants, ultraviolet radiation). There is a relatively well-developed literature studying the effectiveness of EHBE (reviewed in: [4]), which largely parallels the development of the microbiological aspects of hand hygiene. Like hand hygiene, most of the available literature is concerned with the prevention of healthcare-associated infections (HAI) [23–27], with little work done in other settings (but see e.g. [28]). EHBE is much less regulated than hand hygiene, but there have been attempts to regulate and recommend best practices in healthcare settings [29,30]. Concern with EHBE stems from evidence that HCWs can spread infection through the touching of contaminated surfaces [23,31–35], and interventions largely focus on the sterilization of surfaces which might serve as reservoirs of harmful microbes [4].

### Current research on hand hygiene

The common misconception that “all microbes are germs” is apparent in the majority of studies of hygiene, reflected in the focus on bulk reduction in microbial load — even those conducted by clinical microbiologists [36–39]. The concepts of hygiene and sterilization are often conflated, which is perhaps unsurprising given the history of hospital sanitation practices, which seeks to remove all microbes from the environment[16]. There is a logical link between bulk reduction in microbial load and reduction in pathogen spread; however, relatively few studies go beyond cleaning and link hygiene directly to health outcomes, and many of these are specifically concerned with nosocomial infection [37].

Hand hygiene research has focused largely on hospital settings and the spread of nosocomial infection (reviewed in [3,37]), in part due to the history of the field, but moreover because of recognition of the increased risk of infection in places where potentially contagious and immunocompromised people are gathered. Where work on hand hygiene has taken place outside of hospital settings, it has focused on other areas with high risk of pathogen transmission, such as childcare facilities [40–42] or food handling situations [43], or has been undertaken in combination with efforts to improve community environmental hygiene in developing countries (reviewed in [18]).

Between 1980–2001, Aiello and Larson found 53 studies that explicitly linked hygiene to health outcomes outside of healthcare settings, out of thousands of studies matching their search criteria [18]. Studies linking hygiene intervention to health demonstrate the effectiveness of handwashing at reducing the risk of diarrhetic disease [44,45] and upper respiratory infection [45,46]. Reduction in the rates of handwashing in response to fears of lead contamination have been suggested as a factor contributing to a recent *Shigella* outbreak in Flint, Michigan [47].

It is important to recognize that handwashing is just one component of hand hygiene: the role of hand drying as an aspect of hand hygiene has been largely ignored until recently[48]. Recognition of the role that residual moisture plays in the transfer of microbes between surfaces [31,49–51] has focused limited attention on this issue, but there remains no consensus to inform recommendations from regulatory agencies; the WHO *Guidelines* include just three paragraphs on hand drying [3], and note that, “Further studies are needed to issue recommendations on this aspect.”

Most of the existing literature and the prevailing understanding of hygiene is based on cultivation-dependent studies (Fig. 1A), which entail the growth and enumeration of bacteria in the laboratory. These techniques fail to account for the high abundance and ubiquity of non-harmful — and potentially helpful — bacteria on human skin [52,53]. Modern cultivation-*independent* techniques (Fig. 1B), including high-throughput DNA sequencing technology, have facilitated a deeper exploration of microbial diversity and expanded our understanding of the trillions of bacteria, fungi, and viruses living on the healthy human body, collectively known as the **microbiota**, and their role in maintaining health [54]. Studies that focus on hygiene should take this diversity into account and recognize that not all microbes are harmful, and that there is a *continuum* between pathogenic and commensal microbes. Despite the growing use of these modern sequencing technologies, there have been no cultivation-independent studies investigating the *direct* effect of hand hygiene and/or product use on the hand microbiota [55].

**Figure 1:**
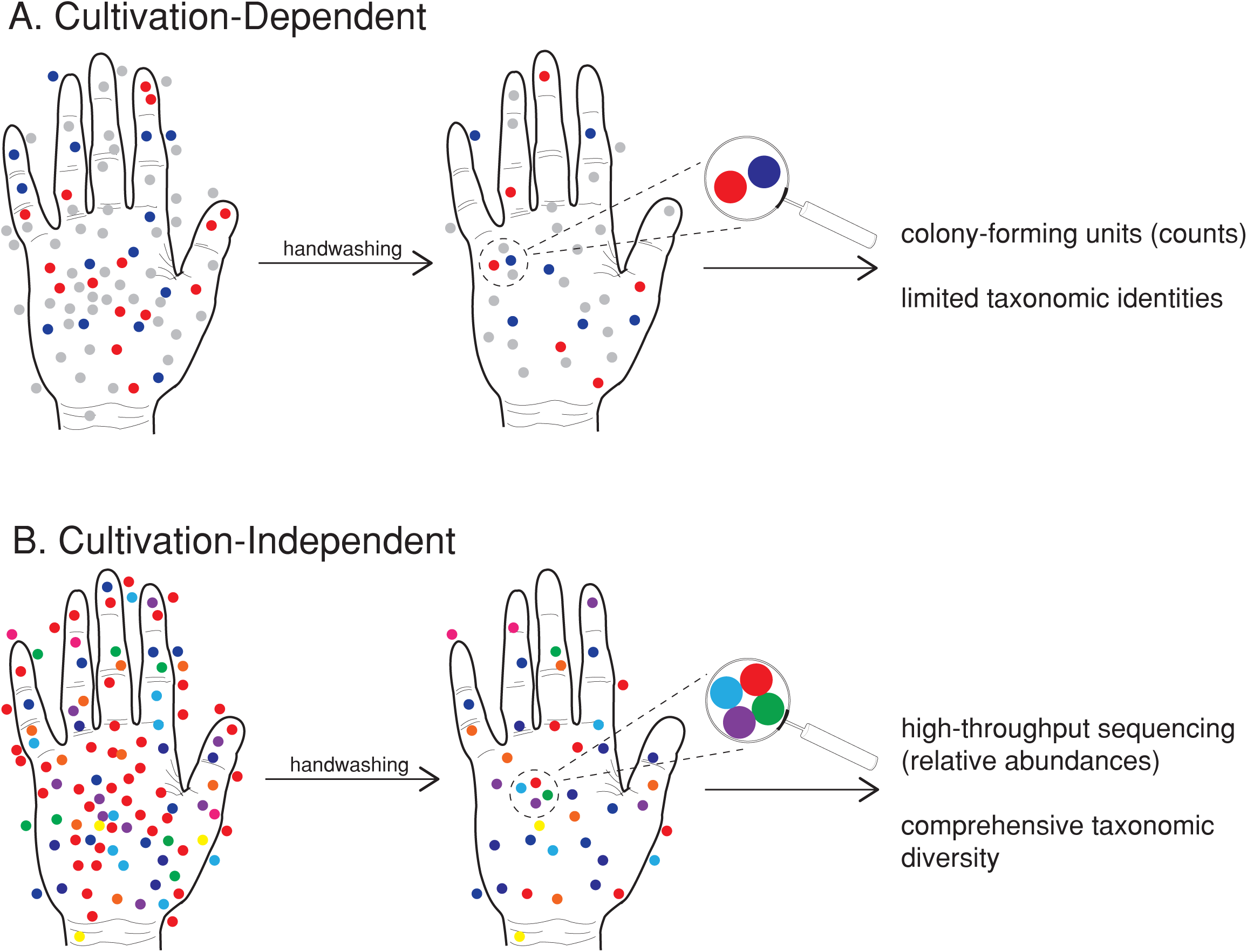
Cultivation-dependent methods. **(A)** are commonly used to study aspects of hand hygiene; many microbes are not detectable using this methodology (represented in grey). Handwashing reduces bulk microbial load, and cultivation yields data showing changes in the numbers of colony-forming units (counts); some studies identify colonies using morphological or molecular methods, yielding limited taxonomic information. **Cultivation-independent methods (B)**, including high-throughput DNA sequencing, are commonly used to study the microbial ecology of the skin. Using these methods, it is possible to quantify alterations in relative abundance of bacterial populations with treatment (such as handwashing), obtain deep, comprehensive taxonomic diversity estimates; depending on technique, it may be possible to also obtain information on functional metabolic pathways (using metagenomics), assessment of proportion of the community that is active (using rRNA / rDNA comparisons, or live/dead cell assays), among other things.

##### Box 1: A note on terminology

Much of the clinical and ecological literature related to the human skin microbiota utilizes two different sets of vocabulary: *resident* and *transient* microbes (often used in the clinical literature; e.g., [56]), versus *commensal* and *pathogenic* microbes (often used in the ecological literature; e.g., [57]). However, these terms are rarely defined and are frequently conflated; residents are often assumed to be commensal, and transients are often assumed to be potential pathogens.

Historically, *resident* microbes were thought of as those that were stable on human skin and were difficult to remove, whereas *transient* microbes were thought to be acquired by contact and could be easily removed from the skin [58]. This notion of resident/transient microbes has continued for decades and has morphed into the assumption that resident microbes are those that commonly reside on skin whereas transient microbes are viewed as contaminants [59].

*Commensal* microbes on human skin are regarded as those that are not typically associated with disease [60]. However, the ecological definition of commensalism refers to the condition where only one organism receives benefit and the other organism suffers no harm [60].

The use of the term “commensal” to describe non-harmful microbes on the skin is suggestive that only the microbe is receiving benefit from living on the skin’s surface and no benefit is provided to the human. However, this definition is misleading because there is growing evidence that microbes once thought to be commensal may actually be involved in host defense, which would suggest a *mutualistic*, rather than a commensalistic, relationship [60]. In comparison, a *pathogenic* microbe on the skin is one that causes harm to the host. There are, however, many microbes that are associated with disease which exist as normal members of the skin microbiota in healthy individuals. When the ecological relationship between host and microbe is unclear (that is, when it is impossible to say if a given microbe is acting as a mutualist, a commensal, or a pathogen), we prefer the term *symbiont* (literally: “together living”), which does not imply an ecological mode.

Both of these dichotomies represent **continua**, which are related, but orthogonal to each other. The idea of a mutualist–pathogen continuum has been successfully applied in the plant microbial ecology literature for decades [61,62]. This continuum represents a position in niche space, which can change through alterations to microbial or host genetics, environmental conditions, and community context [60,63–65]. The resident–transient continuum represents a temporal dimension, and is defined by the length of time that a given microbe is associated with its host — though we must consider the effect of the limits of detection with current techniques [66]. It is important to recognize that resident does not necessarily equate to commensal, nor does transient always mean pathogenic.

It is very well possible for human skin to have mutualistic, commensal, and pathogenic microbes as part of its resident “core” microbiota; a single microbial species may be all of these things. For example, the bacterium *Staphylococcus epidermidis* is commonly found on human skin and is generally regarded as commensal [60], although it can occasionally act as a pathogen [63,64] or a protective mutualist [67]. Recent evidence suggests that *S. aureus*, which has been typically thought of as a pathogenic microbe, is commonly present on healthy skin, specifically in the nasal area [68]. Following this logic, it is likely that the transient microbes people are exposed to in the environment are not only non-pathogenic, but in fact could be beneficial to skin microbiota.

## The human-associated microbiota

The skin is the largest organ of the human body in terms of surface area (1.8 m^2^), serving as the interface between our bodies and the environment [64]. It is home to a vast array of microorganisms — up to 10^7^ bacteria per square centimeter — as well as a great diversity of archaea, viruses, fungi, and mites, generally termed the **skin microbiota** (Box 1) [69,70]. Resident skin microbiota have evolved to take advantage of the relatively harsh environmental conditions found on the skin; in general, skin habitats can be divided into three broad groups: sebaceous, dry, and moist physiological environments. The skin microbiota is far less well studied than the gut microbiota, and the non-bacterial inhabitants of the skin (fungi, viruses, archaea, eukaryotes & protists) are even less well characterized, in large part due to methodological issues, perceived rarity, and asymptomatic nature [71,72].

### Skin habitat and microbial diversity

Human skin may be open to colonization from the environment, but it is thought to be a strong selective filter, largely unsuitable for most microbes to permanently reside [73]. The three major skin habitats (sebaceous, dry, moist), and the gradations of environmental conditions within and between them, largely determine the bacterial community living at a particular skin site [69,74]. Skin bacterial communities, therefore, appear to have **generally predictable biogeographic patterns**. The normal/healthy skin microbiota is composed of a limited number of types of bacterial species (mainly Gram-positive species) [64,73–76]. Dry regions, such as the forearm and palm, are often the richest in bacterial diversity, generally less restricted in membership, and are more susceptible to temporal variability, while sebaceous sites are generally poorer in bacterial diversity and dominated by *Propionibacterium acnes*, presumably due to high sebaceous gland activity that may result in more exclusivity [64,74,76,77].

While there is some similarity in microbiota of similar body habitats and across individuals, it is abundantly clear that not all skin communities are alike [64,66,74,76,78,79]. Although only a small fraction of certain taxa found within a single body site have been detected at every sampling time point for up to a year, many taxa have still been identified as persistent community members if they appeared in a given body habitat for an extended period of time [56,66]. Nevertheless, it is difficult to define a **core microbiota** for a given anatomic site on the skin [80]. A recent review listed the most “common” human skin bacterial residents as: *Staphylococcus, Corynebacterium, Propionibacterium, Micrococcus, Streptococcus, Brevibacterium, Acinetobacterium*, and *Pseudomonas* [60]. Many of these (e.g., *Staphylococcus aureus* and *S. epidermidis*) also have the potential to become multidrug resistant pathogens, emphasizing the insidious nature of classifying microorganisms as one ecological mode (e.g. commensal v. pathogenic) and the need for a conceptual framework taking into account the existing ecological continua (Box 2).

##### Box 2: Ecological context and the skin

There is an emerging appreciation of the microbial ecology of the skin. *Community ecology* seeks to understand what factors determine the presence, abundance, and diversity of species in a community [81]. Island biogeography theory [82], in particular, allows us to conceptualize each person as an island: a patch of habitat that must emerge and assemble its communities by the fundamental processes of community ecology. The interactions between skin microbial communities and the host makes understanding the ecological factors contributing to microbial communities particularly important. Multiple ecological factors interact to determine the species composition in a given ecological community; *dispersal* (Fig. 2a) and *environmental selection* (Fig. 2d) are the two factors most relevant to the discussion of hygiene.

**Figure 2:**
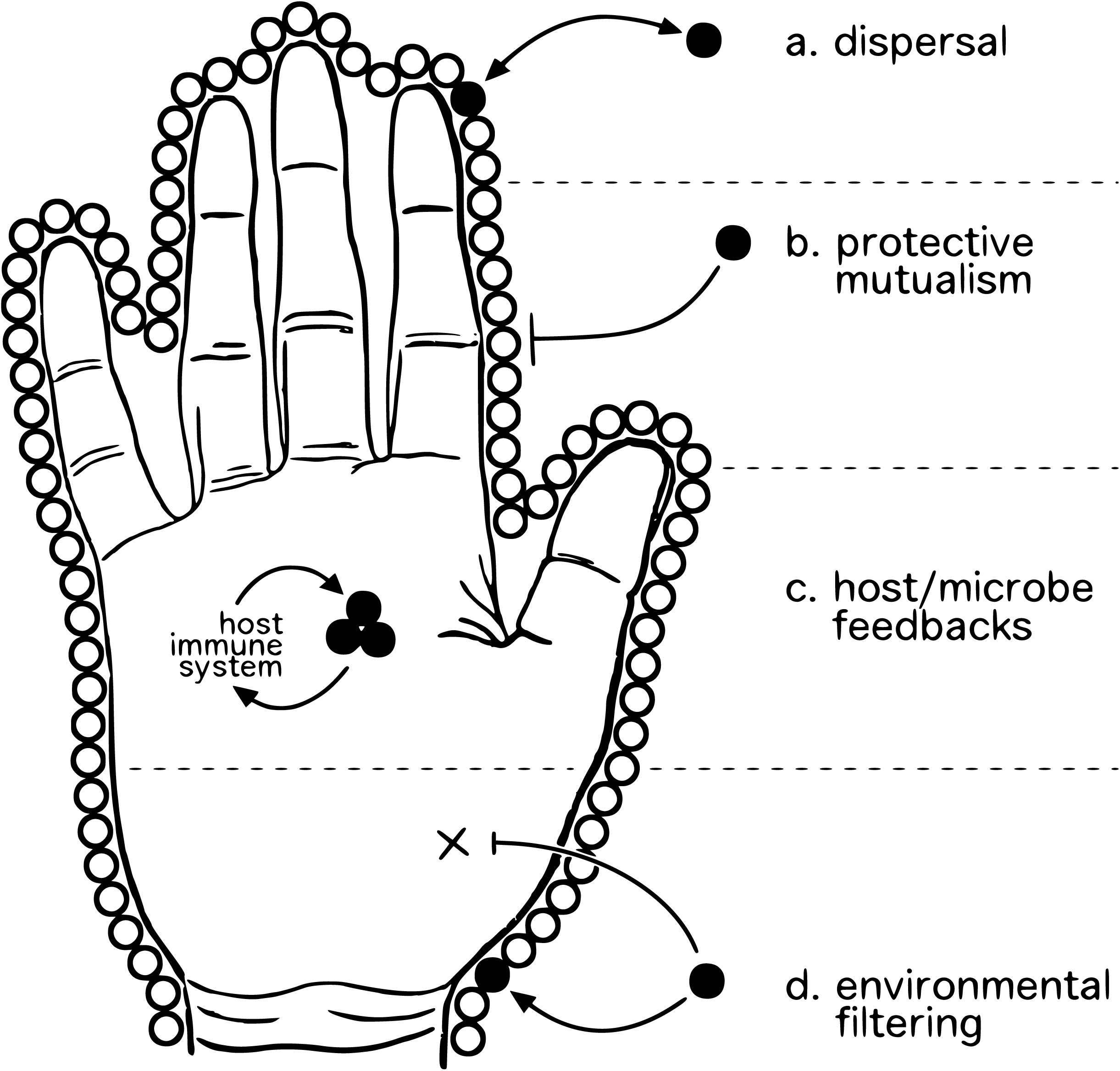
Conceptual illustration of important ecological factors impacted by hygienic practice. *Dispersal* (a) is the movement of organisms across space; a patch of habitat is continuously sampling the pool of available colonists, which vary across a variety of traits (dispersal efficiency, rate of establishment, *ex host* survivability, etc.) [93]; high dispersal rates due to human behaviors (e.g., microbial resuspension due to drying hands with an air dryer) have the potential to disperse both beneficial and harmful bacteria alike. *Protective mutualisms* (b) function through the occupation of niche space; harmful microorganisms are excluded from colonization via saturation of available habitat by benign, non-harmful microbes [94]. *Host/microbe feedbacks* (c) occur via the microbiota’s ability to activate host immune response, and the host immune system’s ability to modulate the skin microbiota [95–97] — multiple pathways, including IL-1 signaling [67] and differential T-cell activation [98], are involved — such feedbacks between host immune response and the skin microbiota are thought to be important to the maintenance of a healthy microbiota and the exclusion of invasive pathogenic microbes [99]. *Environmental filtering* (d) works on the traits of dispersed colonists — microbes that can survive in a given set of environmental conditions are filtered from the pool of potential colonists [93]: the resources and conditions found there permit the survival/growth of some organisms but not others. The importance of *diversity* of the microbiota to each of these ecological factors should not be underestimated; interactions between taxa may modulate their ecological roles, and community variation across a range of ecological traits may be altered by changes in community membership or structure [100].

**Dispersal** (Fig. 2a) of commensal or mutualistic organisms may be particularly relevant to human health. Studies that have examined the transmission of human-associated microorganisms have almost exclusively focused on pathogenic microbes in healthcare settings [81]. The transmission of other (i.e., non-pathogenic) members of the skin microbiota is poorly understood, including the roles of a number of factors (e.g., diversity, interspecies interactions, host factors, environmental factors) on the ease of microbial transfer and subsequent colonization. Transmission via direct contact with other individuals, or indirectly with fomites or water droplets, introduces transient microbes that could alter the ecological dynamics of the skin microbiota [81].

All persons are dispersers of their microbiota, though dispersal rates vary within and among people. Any organism living on the skin will be dispersed as a result of normal desquamation (i.e., shedding or peeling of the outermost layer of skin) [83]. Individuals emit a personalized microbial cloud that likely impacts both coinhabitants and the microbiota of the built environment itself [84]. While research on whether resident microorganisms can be transferred among individuals is nascent, it is hypothesized that delivery method at birth (vaginal vs. Cesarean section) affects initial skin microbial communities of infants [85] (but see [86]).

**Environmental filtering** (Fig. 2d) of dispersed microbes functions primarily through differences in skin habitat (e.g., dry vs. sebaceous sites). Interactions between microbial populations may be part of the “filtering” of the environment; thus, *priority effects* and factors related to the established microbiota can be considered part of the environment to newly dispersed microbes. There is evidence that host factors vary in the ability to promote bacterial colonization, and that this varies by skin site [76]. The role of *host/microbe feedbacks* (Fig. 2c) in determining environmental selective pressures may also influence the outcomes of potential dispersal events. There is some evidence that microbial communities may be transferred between people or their environments [87,88].

*Invasion ecology* focuses on perturbations of established communities, and attempts to understand the factors that allow invasion by exogenous species [89,90]. Applied to the skin microbiota, disturbance (e.g., hand hygiene practices) may be a major factor in alterations of the skin microbiota through invasion; *protective mutualisms* (Fig. 2b) may be disrupted or eliminated, allowing invaders to colonize. The frequency and magnitude of such disturbances likely facilitate the invasion of potentially undesirable, pathogenic species [91,92].

A conceptual framework for understanding the interactions between skin microbiota, the human host and environment, and the impact on human health must take into account all of these ecological factors [81,91]. Significant and potentially harmful alterations of the skin microbial community structure may occur as a result of several factors: dispersal of non-resident microbes to the host microbiota, disturbance regimes (e.g., handwashing practices), local and regional environmental factors (i.e. environmental selective filters on source and sink populations, such as host skin condition and indoor settings), and the genetics and demographic characteristics of the host, which also provide selective filtering [81].

### Skin microbiota function & role in host immunity

The resident microbiota has evolved in conjunction with the human host and is thought to be important to the maintenance of healthy ‘normal’ skin function. Generally, the resident microbiota have a positive effect on human health through protective mutualism (Fig. 2b); it is only when the host becomes compromised that the resident microbiota displays pathogenic potential [99,101]. As the skin is our body’s interface with the outside world, it must act to both prevent colonization by pathogens and tolerate or encourage the presence of potentially protective bacteria. The skin is a complex immunological organ with both innate and adaptive immune cells, including multiple dendritic and T-cell subsets; antimicrobial peptides, proinflammatory cytokines, and chemokines that are secreted by keratinocytes to support an immune response [102–106]. While pathways related to infection response are relatively well-understood [60,77,99], the mechanisms by which commensal or transient bacteria are tolerated by the cutaneous immune system are less well known.

Host/microbe feedbacks (Fig. 2c), modulated through the host immune system, have been recently demonstrated, and are likely to play critical roles in maintaining healthy host/microbiota relationships. For example, *Staphylococcus epidermidis* has been shown to produce antimicrobial peptides and may modulate the host immune response [60]; *S. epidermidis* and *Corynebacterium* spp. are capable of reversing or preventing the successful colonization of *S. aureus* in the human nares [71], such that removal of *S. epidermidis* may be harmful to the host through increased colonization of pathogens [60]. Skin dysbiosis has been linked to many skin disorders, including acne vulgaris, psoriasis, and atopic dermatitis [107–114]. Investigation of the potential of microbiota transplants and probiotic skin treatments for these diseases are underway [64]. Thus, **skin microbiota is likely of direct benefit to the host, and only rarely exhibits pathogenicity**.

### Disturbance

The microbial ecosystem of the skin is relatively stable in the face of continual desquamation of the skin surface and frequent perturbations [79]. There is a lack of research on the role of skin microbial communities, and disturbance of those communities, on the risk of infectious disease transmission [81]. Disruptions by antibiotics, handwashing, or cosmetic application may alter the microbial community, enabling invasion of pathogenic microbes or a shift in dominance leading to dysbiosis [81]. Skin disturbance can predispose the host to a number of cutaneous infections and inflammatory conditions [60]. For example, *S. aureus* — once believed to be a transient colonizer — is an extremely common member of the human skin microbiota that somehow turns pathogenic upon disturbance [115]. Handwashing is a frequent disturbance of the skin microbiota, possibly perturbing the existing trade-off between its microbial colonizers and competitors. Despite the multitude of studies emphasizing the benefits of personal hygiene on reducing disease transmission by removing or reducing transient microorganisms, the effects of handwashing on the *resident* microbiota are not well studied. Different behavioral habits (e.g., frequency and duration of washes, product used, etc.), likely account for at least some of the variation in microbial community structure and membership observed in human studies [81].

### Hand hygiene and the microbiota

Hands harbor greater bacterial diversity and are more temporally dynamic than other body sites [55]. More than 150 bacterial species have been recovered from human hands; these species primarily belong to the phyla Firmicutes, Actinobacteria, Proteobacteria, and Bacteroidetes [55,75]. This increased diversity on human hands compared to other skin sites may be a result of the exposure of hands to consistently varying external environments.

Like other skin sites, there is a high degree of interpersonal variation in the hand microbiota; a minority of taxa (13%) are shared between the hands of any two individuals, and the two hands of a single person may share only a slightly larger fraction, though those communities appear somewhat stable [75]. Additionally, the temporal dynamics of the hand microbiota are unpredictable [66,116]. Despite the evidence that certain bacterial species remain present on the hands over time, their relative abundances are variable [66]. Both internal and external factors may also modulate the microbial communities on hands [55,64]. Microbial communities on people’s hands are significantly affected by host factors, including sex, relatedness, living quarters, hand hygiene, and even pet ownership [55,75,117].

Hand hygiene is still regarded as the most important practice to prevent the transmission of microbes and minimize the spread of disease [118]. However, compliance with hand hygiene practices in healthcare settings is generally low, with mean baseline rates ranging from 5 to 89% [3]; typical rates may be no better than 40% [16,118].

Current understanding of the effects of hand hygiene in healthcare settings largely stems from cultivation-based methods focusing on identification of pathogenic microbes. These clinical studies have historically been performed during periods of infectious outbreak in hospital settings with the assumption that bacteria on skin are pathogenic contaminants [119]. Even with the growing use of high-throughput sequencing, there have been no cultivation-independent studies that have investigated the direct effects of hand hygiene or product use on the hand microbiota [55]. There is great potential to further our understanding of the human hand microbiota by utilizing an ecological perspective in healthcare settings, where hygiene practices are vital. Despite this current gap in knowledge, we are still able to draw preliminary conclusions about hand hygiene and its effect on the skin microbiota from cultivation-based studies and the few cultivation-independent studies that have looked at this relationship indirectly.

In cultivation-based studies, the length of direct patient contact is positively correlated with bacterial counts [118], and surface area and time of contact significantly affects the abundance of bacteria present on the hands of HCWs [120]. Older work has shown that soap and water handwashing is effective at removal of patient-acquired microbes [121], and more recent studies have shown alcohol-based handrubs to be as effective [118,122] or even superior to soap and water [59]. There is also an interaction between skin health and the effect of hand hygiene that may be of concern: increased handwashing may increase the amount of microbes on hands due to worsening skin health [123]. Additionally, moisture level has a significant effect on cross-contamination rates [31,49–51]. However, these studies examined bacterial load on hands and failed to address the *identities* of the species that were affected by hand hygiene practices — identity matters when most members of the microbiota are commensal or even potentially mutualistic.

Cultivation-independent studies show some similar trends: the use of alcohol-based products for hand hygiene may significantly reduce bacterial diversity on HCWs’ hands; however, this decreased bacterial diversity may increase the likelihood of carrying potential pathogens on the hands by eliminating naturally occurring protective or commensal species [124]. Time since last handwashing was significantly correlated with changes in bacterial community composition but did not affect bacterial diversity [75]. This result could suggest that the bacterial community present on hands quickly reestablishes itself post-handwashing, or that not a lot of bacterial taxa are removed during the handwashing process [75].

In order for human-to-human microbial transmission to occur in a healthcare setting, microbes must be transferred from a patient to a HCW, the microbe must be capable of surviving for a period of time and hand hygiene must be inadequate to remove the microbe, and the HCW’s contaminated hands must come into direct contact with another patient [125]. One study that looked at the transmission of *Klebsiella* spp. among HCWs in an intensive care unit found that only slight contact with patients was needed to transfer the microbe to HCWs and that *Klebsiella* spp. could survive on dry hands for up to 150 minutes [126]. Another study found that the transmission potential of microbes to and from hands and sterile fabrics was highly species dependent, suggesting that hygienic practices may play a more vital role in transmission prevention of certain microbes over others [122].

More studies are needed to quantify the role of interactions with the resident hand microbiota in the transmission of potentially pathogenic microorganisms. From the available data, we can conclude that transmission via hands is common and often related to microbial load, and that variation in moisture levels affects transmission efficiency.

### Human-associated microbes and the built environment

As research on the human microbiota continues to expand and diversify, many researchers have turned their attention to the microbiology of the indoor spaces we inhabit for the majority of our lives: the built environment. Until recently, microbes indoors have largely been viewed in a negative light as potential pathogens that may cause illness if transmitted to humans via surface contact, inhalation of bioaerosols, or through altered patterns of human-to-human interactions influenced by the built environment. This perspective is reflected in the near-ubiquity of antimicrobial chemicals in typical household cleaning products, along with increasing use in building materials and other consumer goods [128]. Despite efforts to sanitize the built environment, research shows that the indoors is nevertheless teeming with microbial life. The ways in which we design, operate, and occupy buildings can have significant impacts on the kinds of microbes that reside in our homes, hospitals, offices, and public dwellings.

Numerous studies have reported that humans are predominant contributors to microbial communities found indoors. In addition, pets, houseplants, and even the fruits and vegetables we bring inside carry their own microbes with them (reviewed in [129–131]). Abiotic factors such as plumbing and ventilation systems also play a significant role in shaping the indoor microbiota, and studies in hospitals [132] and university office and classroom buildings [117,133] demonstrate that natural ventilation introduces outdoor microbial communities and can diminish the human microbial signature. Deposition of microbes occurs rapidly after humans move into a space [134] or when natural ventilation is flushed through a building [117]. Shortly after disinfection of surfaces, microbial communities in athletic gyms [135] and public bathrooms [136,137] are quickly reestablished even in the absence of direct human contact or occupancy. In environments such as hospital neonatal intensive care units (NICUs), where removal of potential pathogens is seen as critical to infant health, disinfection practices do not remove all microbes or reduce microbial diversity on surfaces [138].

Given these findings, we can begin to view microbes indoors not as primarily pathogens, but rather as communities of microbes that are common residents co-inhabiting our indoor spaces with us. Indeed, the *hygiene hypothesis* is gaining support, and some studies have shown that microbial exposure—particularly early in life—may have long-term implications for health and immunity [139,140]. If the microbes indoors are an important component to this microbial exposure, it behooves us to adopt a broader perspective of indoor microbial ecology. This parallels our evolving perspective of the human skin microbiota. Likewise, our understanding of best practices for environmental hygiene of the built environment are likely to evolve in the near future as our understanding of the microbiota of the built environment changes. Indeed, there is already suggestion that environmental probiotics may be effective at reducing the spread of pathogenic microbes in hospitals [141,142].

## Defining hygiene

Understanding the ecological dynamics within human-associated microbial communities gives us the power to improve strategies for the maintenance of our microbiota for health and informed management of the crucial health-associated ecosystem services provided by these microbial communities. If the desired outcome of hygienic activities is to improve health, and health is improved through optimal microbial maintenance and management within the host, then we would do well to have hygienic guidelines that bear this in mind.

The evidence that microbes are essential for maintaining a healthy skin microbiota supports the idea that hygienic practices aimed at the simple removal of microbes may not be the best approach. Rather, hygienic practices should aim to reduce *pathogenic* microorganisms and simultaneously increase and maintain the presence of *commensal* microorganisms essential for host protection. It is clear that microbial colonization of the skin is not deleterious, *per se.* Humans are covered in an imperceptible skim of microbial life at all times, with which we interact constantly. We posit that the conception of hygiene as a unilateral reduction or removal of microbial load has outlived its usefulness and that a definition of hygiene that is quantitative, uses modern molecular biology tools, and is focused on disease reduction is needed. As such, we explicitly define hygiene as **‘those actions and practices that reduce the spread or transmission of pathogenic microorganisms, and thus reduce the incidence of disease’**. To examine the implications of thinking about hygiene in this way, we examine one aspect of the hand hygiene literature in some depth: hand drying.

### Hand drying and hygienic efficacy

There has been some controversy in recent years concerning the role of hand drying in hand hygiene practices. Most sources agree that it is of great importance [3,16], but there has been comparatively little work done to quantify the contribution that drying makes to pathogen transmission. Early hand hygiene studies found that handwashing with soap and water was much more effective at removing bacteria when hands were dried with a paper towel then when hands were allowed to air dry [143]. Other work supported this, showing increased bacterial transmission from improperly dried hands [50,144]. Patrick and colleagues [49] explicitly tested the relationship between residual moisture on the hands and bacterial transmission, finding that wet hands were much more likely to transfer bacteria between objects. Despite recognition of the importance of drying to overall effectiveness of hand hygiene and potential impact on infection control [31,49–51], little effort has been made to date to quantify the impacts of hand drying regimes [38,48].

Because of the general lack of free water on the skin, localized changes in hydration have a significant effect on microbial load. Naturally occluded sites have high densities of colonizing microbes compared to areas with low free water [145]. There is some evidence that microbial species are differentially affected by water availability; some thrive in wetter conditions on the skin than others [101]. Effects of residual moisture on hand hygiene may then be two-fold, both by increasing transmissibility [49,50,143,144], and by altering community composition through environmental filtering [81,145].

Much of the existing work on hand drying has examined the “hygienic efficacy” of various methods — typically paper towels, warm air dryers, and jet air dryers. What is meant by “hygienic efficacy” is often left unstated, but usually is measured by change in microbial load, dispersal of microbes from the hands, or some proxy thereof.

*Warm air driers* are those that blow a heated stream of air, generally for 30–40 seconds per use, under which the user typically rubs their hands together during drying. Most research has shown that warm air dryers may increase the number of bacteria on the hands after use [144,146–149], with some exceptions showing no change [148,150–154] or a reduction [155–157]. This increase in bacterial counts could be the result of the existing bacteria within the dryer mechanism [146,149], the re-circulation of microbe-enriched air [129,158] (including an enrichment of fecal-associated bacteria [144,149]), the liberation of resident bacteria from deeper layers of the skin through hand rubbing while drying [38,147,157], or some combination of the above. It is also important to note that the temperature of warm air dryers is not hot enough to kill bacteria [151]; the purpose of heating the air is solely to aid in evaporation. Additionally, warm air dryers are slower at drying the hands [38,39,48,144
146,–149,158–160], which is thought to reduce compliance with drying (i.e., people walk away with wet hands).

A recent alternative to warm air dryers are *jet air dryers*, which typically use a high-speed jet of unheated air to push water from the hands, typically achieving drying in 10–15 seconds. Research on *jet air dryers* has focused on the importance of the total dryness of hands, contrasting the speed of jet air drying with that of warm air dryers and emphasizing the risk of cross-contamination with wet hands [38,39,153,160]. These studies typically employ cultivation and enumeration to measure the number of bacteria transferred and use residual moisture to measure efficiency of drying. The reduced drying times achieved by jet air dryers are noted repeatedly [39,161,162], with drying times that are generally comparable to paper towels [39,48]. Many jet air dryers (e.g., the Dyson Airblade^TM^) are marketed as designed with a high-efficiency particulate air (HEPA) filter built into the airflow system, which reduces the risk of redistribution of airborne microbes to the hands [38]. However, there is concern about the propensity of such rapid air movement to aerosolize microbes from users’ hands or the surrounding environment, as evidenced by the number of studies examining the dispersal of microbial suspensions or some proxy thereof by such devices [39,153,160,161,163]. Particular attention has been paid to the distance such rapid air movement is capable of dispersing potentially contaminated droplets from the hands, though methods typically employed unrealistic microbial loads, or artificial proxies such as paint [39,160,161,163].

Drying with *paper towels* is the method recommended for healthcare workers by both the Centers for Disease Control and Prevention [17] and the WHO [3], due in large part to bulk bacterial count data indicating that paper towels are effective at removing surface bacteria [48,144,147,149,157,160,164]. Use of paper towels is also associated with only minimal spread of droplets from the hands into the environment [39,160,161,163,165,166], though it is possible that waste paper towels may serve as a bacterial reservoir [150,153]. Additionally, there is great variance in the manufacture and storage of paper towels, which may lead to risk of contamination as part of the manufacturing process, particularly of recycled paper towels [166].

*Cloth towels* represent a final alternative, though they are seldom used in modern public facilities due to risks of potential cross-contamination at the end of the roll [48,167]. This is despite some evidence that roller-type towels are highly efficient at drying the hands, unlikely to be contaminated (likely due to the steam cleaning process), and comparable to paper towels at bacteria reduction, provided each user has access to a fresh section of the roll [151].

Several Life Cycle Analyses (LCA) have compared other aspects of these different drying systems, including cost effectiveness and environmental impacts [168,169]. In general, impacts are driven by usage, rather than manufacturing or maintenance, and paper towels tend to have greater environmental impacts because the energy costs inherent in shipping bulky materials outweighs the energy necessary to run most air dryers. A holistic consideration of environmental impact of hand drying would include efficacy according to the definition of hygiene we have offered, which may be more important in some contexts than others (such as hospitals).

### Recontextualizing cleanliness in hand drying

The hand drying literature can be separated into two opposing divisions: one attempting to demonstrate that the newer air dryers are as hygienically efficacious as paper towels [38,150,155,166], and the other attempting to discredit the newer technology in favor of paper towels [39,48,146,160,161,163,164]. While both divisions utilize bulk reduction in microbial load as a proxy for hand hygiene [38,160], research from the first division largely focuses on the potential of wet hands to transfer microbes [49] and the ability of air dryers (whether warm or jet) to effectively dry hands [38,39,150,153]: the hypothesis in this case is drying is hygienically efficacious *if hands are dry* and new microbes are not acquired through the process. Research from the second division tends to focus on the risk of air dryers to spread microbes throughout the environment by aerosolizing moisture from the hands [39,160,161,163,165]: the hypothesis in this case is drying is hygienically efficacious if new microbes are not acquired through the process and *if production of aerosols are minimized.* It is difficult to compare the two divisions because many of these studies include methodological issues (e.g., variation in protocols, lack of appropriate controls or statistical analyses) that make it difficult to compare results across studies.

Despite there being an obvious interplay between these two divisions, many of the concerns on either side remain unaddressed. **Utilizing a definition of hygiene that explicitly relies on reduction in disease spread** would address concerns on both sides of the debate: there is currently no evidence linking aerosolization of residual moisture (and associated microbes) with the actual spread of disease. Likewise, despite demonstrations that wet hands allow for increased bacterial transmission, there does not seem to be evidence linking wet hands after washing to deleterious health outcomes. The complex ecological context of the hand microbiota may modulate effects of both aerosolization and prolonged moistening. Additionally, the majority of hand drying research largely ignores the relative hygienic contribution of the hand *washing* step [38,150,160,161,163]; understanding the relative contribution of washing to hygienic efficacy is necessary to put the hand drying literature in proper context.

Future research should take advantage of cultivation-independent techniques (Fig. 1), explicitly include the contribution of handwashing (and other controls necessary to accurately interpret results) and work to increase sample size to ensure statistical rigor. Such research should work to bridge the gap between the two existing divisions of research by using health outcomes (such as the spread of disease) as dependent variables, taking into account the microbial community context of the skin microbiota, and focusing on understanding the relative contribution of bioaerosols and residual moisture to the risk of disease transmission. Working to link the effects of human behaviors, such as dryer usage, to the microbiota of the built environment will help to link our understanding of hand hygiene and environmental hygiene of the built environment.

## Conclusions

Concepts of hygiene have evolved greatly over the last few centuries, influenced by cultural norms of cleanliness, empirical data, and the advent of the germ theory of disease. Through widespread acceptance of the germ theory, the common misconception that “all microbes are germs” has come to influence the modern usage of *hygiene*, such that it has become nearly synonymous with *sterilization.* The history of regulation of hygiene in healthcare related settings generally reflects this usage. Modern microbial ecology using sensitive, cultivation-independent techniques provides a glimpse into the complexity of the microbial communities in, on, and around us, as well as a growing appreciation for the ecosystem services provided by these microbial communities.

Given the intimate interactions between humans and our microbiota, it is becoming apparent that maintenance and promotion of healthy human-associated microbial communities is necessary for the maintenance of good health. As such, we argue that the concept of hygiene as akin to sterilization no longer serves a useful role in scientific or medical discourse. It is more useful to explicitly define hygiene in terms of health outcomes, and focus on the use of quantitative, modern molecular biology tools to elucidate the complex ecological interactions that relate hygienic practice to the spread of disease. Pursuant to that goal, we have explicitly defined hygiene as ‘**those actions and practices that reduce the spread or transmission of pathogenic microorganisms, and thus reduce the incidence of disease**’.

Using such a definition alters the way we approach research on hygiene, and suggests novel avenues of research. Studies of skin dysbioses [60,77,107] are beginning to demonstrate that consideration of species identity & ecological context is necessary to understand disease progression and devise effective treatments in some cases. Consideration of microbial ecological context as it relates to hygienic practice may improve understanding and treatment of many skin diseases, including atopic dermatitis, psoriasis, and acne. Already, methods similar to the gut microbe transplantation used to successfully combat *C. difficile* infection are now under consideration for common skin diseases [64].

To our knowledge, no studies have used the drying step of hand hygiene protocols as an independent variable and *health outcomes*, such as disease transmission or development of symptoms, as a dependent variable. Indeed, very few studies of hand hygiene examine health outcomes as a response at all [18]. Nearly all studies of hand drying utilize bulk reduction in bacterial load as a proxy for reduced transmission of pathogenic organisms [48]. However, due to the complex microbial ecology of the skin [76] and the potentially differential effects of such disturbances have on different microbial species [81], such a proxy is likely to not be broadly appropriate: it is necessary to know the identities and ecological roles of the organisms affected. New methods — including those that enable the assignment of functional groups to classes of microbes based on cultivation-independent, high-throughput DNA barcode surveys; quantification of the metabolically active portions of microbial communities and live/dead microbial determination methods; and high-throughput, whole-genome metagenomic sequencing, which enables the quantification and assignment of true functional potential — will help us to understand the ecological effects of hand hygiene practices, including hand drying. Explicit quantification of the effects of various hygienic practices on health metrics will allow us to understand the complex interplay between microbial community dynamics, hygienic practices, and health outcomes, and hopefully provide meaningful data to support future recommendations and regulations for hygiene practices.

#### Box 3: Glossary

*biogeography*: the discipline studying the distribution of species and ecosystems in space and across evolutionarily meaningful timescales
*built environment*: artificially constructed and maintained environments; the “indoors”.
*community ecology*: the discipline studying the organization and function of ecological communities (those organisms actually or potentially interacting, bounded by either geographic or conceptual limits).
*contamination*: incidental presence of microbes; not long-term residents of the microbial ecosystem in question.
*cultivation-dependent techniques*: microbiological techniques that rely on the cultivation of microbes for enumeration and identification; less than 1% of microbes are estimated to have been cultivated in the lab [52,53], leaving a vast majority of microbial diversity underexplored (Fig. 2A).
*cultivation-independent techniques*: techniques for the elucidation of microbial communities that do not rely on cultivation of microorganisms; these generally rely on high-throughput, next-generation sequencing technologies (e.g., Illumina, 454 pyrosequencing) that allow for the direct sequencing of DNA or RNA from the environment; common techniques include **metabarcoding**, in which a conserved “barcode” region of the genome is amplified and sequenced from environmental samples, giving information about which taxa are present and their relative abundances, and **metagenomics**, in which all available microbial DNA is sequenced, giving information about presence and relative abundances of metabolic pathways as well as identities of microbes (Fig. 2B).
*dispersal*: the distribution of propagules across space.
*dysbiosis*: perturbation of human-associated microbial communities, such that some members shift to a pathogenic ecological mode.
*environmental filtering*: the process by which potential colonists are selected based on purely ecological factors.
*ecological niche*: a broad term encompassing multiple definitions used to describe to an organism’s activity or behavior in response to a given set of biotic and abiotic environmental conditions or resources. Organisms occupy niches by carrying out specific functions, often through competitive or mutualistic interactions. **Niche space** refers to the set of all possible niches, occupied or unoccupied, in a given habitat.
*hygiene*: those actions and practices that reduce the spread or transmission of pathogenic microorganisms, and thus reduce the incidence of disease.
*hygiene hypothesis*: the idea that a lack of early childhood exposure to microorganisms increases susceptibility to allergic diseases by suppressing the natural development of the immune system.
*invasion ecology*: the discipline studying the alterations to ecosystems resulting from introduction and establishment of taxa originating outside of said ecosystem, and the factors allowing some taxa to invade successfully.
*microbial ecology*: the discipline studying the interrelations between microorganisms, including but not limited to community interactions and interactions with the environment.
*microbial load*: the absolute abundance of microbes; commonly estimated using cultivation-dependent techniques through quantitative counts of **colony forming units** (CFUs).
*microbiota*: or **microbiome**, the ecological community of microorganisms (bacteria, archaea, viruses, fungi, mites, etc.) that share our body space; may be subdivided into cohesive groups, such as the skin microbiota, or the gut microbiota.
*nosocomial*: of or relating to hospitals.
*priority effects*: the particular influence that early arriving members of a community have on later arriving members.
*protective mutualism*: a mutualism in which protection from pathogenic organisms is the result of occupation of niche space within the host habitat, excluding colonization by harmful microbes; often conflated with commensalism (see Box 1).
*sterilization*: the removal of all microbes from a surface or object.
*transmission*: dispersal and establishment of microbes between hosts.

## Abbreviations

CFU: colony forming unit
EHBE: environmental hygiene of the built environment
LCA: life cycle analysis
HAI: hospital-associated infection
HCW: healthcare worker

## Declarations

### Ethics approval and consent to participate

Not applicable

### Consent for publication

Not applicable

### Availability of data and material

Not applicable

## Authors’ contributions

RV outlined the manuscript, performed literature review, and wrote the manuscript. AB, KS, MN, and RH performed literature review and assisted in writing the manuscript. RV and MN prepared figures with input from all authors. JG and KVDW secured funding for the research, contributed conceptually to the manuscript, and provided feedback. All authors read and approved the final manuscript.

## Acknowledgements

The authors are grateful to Jeff Kline and Stephanie Luiere for their feedback on an earlier version of the manuscript.

